# Social framing effects in decision making

**DOI:** 10.1101/2021.09.28.462257

**Authors:** Payam Piray, Roshan Cools, Ivan Toni

## Abstract

Human decisions are known to be strongly influenced by the manner in which options are presented, the “framing effect”. Here, we ask whether decision-makers are also influenced by how advice from other knowledgeable agents are framed, a “social framing effect”. For example, do students learn better from a teacher who often frames advice by emphasizing appetitive outcomes, or do they learn better from another teacher who usually emphasizes avoiding options that can be harmful to their progress? We study the computational and neural mechanisms by which framing of advice affect decision-making, social learning, and trust. We found that human participants are more likely to trust and follow an adviser who often uses an appetitive frame for advice compared with another one who often uses an aversive frame. This social framing effect is implemented through a modulation of the integrative abilities of the ventromedial prefrontal cortex. At the time of choice, this region combines information learned via personal experiences of reward with social information, but the combination differs depending on the social framing of advice. Personally-acquired information is weighted more strongly when dealing with an adviser who uses an aversive frame. The findings suggest that social advice is systematically incorporated into our decisions, while being affected by biases similar to those influencing individual value-based learning.

## Introduction

Human decisions are often influenced by information derived from the behavior of other agents (Baldwin and Baird, 2001; Siegal and Varley, 2002; Saxe, 2006; Behrens et al., 2009; Klucharev et al., 2009; Biele et al., 2011; Izuma and Adolphs, 2013; Stolk et al., 2013; Diaconescu et al., 2014; Chung et al., 2015; Reiter et al., 2017; Chib et al., 2018; Konovalov et al., 2018). Previous work has identified when and how decisionmakers learn from knowledgeable agents (Morgan et al., 2012; Heyes, 2016). For instance, decisionmakers weight socially-acquired information over personally-acquired information when the latter is considered uncertain (Morgan et al., 2012; Toelch et al., 2014), and as a function of the trustworthiness that advisers have built through their behavior over time (Behrens et al., 2008; Biele et al., 2011). Human decision-makers are also greatly biased when decision outcomes are described to them in an appetitive or aversive frame, even when those outcomes are financially equivalent (Kahneman and Tversky, 1979; Kahneman et al., 1982; De Martino et al., 2006; Rangel et al., 2008). Accordingly, framing outcomes is used by agents to influence decision-makers (McKenzie and Nelson, 2003; Sher and McKenzie, 2006; Frith and Singer, 2008; Cukier et al., 2021). Here, we ask how framing an advice, a prototypical socially-acquired information, biases the weight allocated to the advice and/or to the adviser by decision-makers, and alters their learning behavior.

Computational algorithms that support learning the trustworthiness of others over time are hypothesized to be similar to those that support learning about consequences of our own actions, even though those algorithms might be implemented by different brain circuits (Behrens et al., 2008, 2009; Nicolle et al., 2012; Ruff and Fehr, 2014). For instance, the computation of socially- and personally-acquired information is implemented by partially segregated circuits centered around the temporal-parietal junction (TPJ), often associated with social learning; and the ventral striatum, involved in reward learning (McClure et al., 2003; O’Doherty et al., 2003; Saxe, 2006). The ventromedial prefrontal cortex (vmPFC), a region implicated in value-based decision making, has been involved in combining socially- and personally-acquired information, at the time of choice (Behrens et al., 2008, 2009; Ruff and Fehr, 2014). However, it remains unclear how the combination of personally- and socially-acquired information is influenced by advice offered in appetitive or aversive frames, in particular when the reliability of information is uncertain. One possibility is that appetitively- or aversively-framed advice influence the weight allocated to the advice over personally-acquired probabilistic information, thus modulating reliance on the advice at the time of choice. The vmPFC is responsive to both appetitive and aversive values across different modalities (Pessiglione and Delgado, 2015), and it combines personally-with socially-acquired information about the relative value of options when implementing choices (Nicolle et al., 2012; Zaki et al., 2014). Another possibility, not mutually exclusive, is that appetitively or aversively-framed advice influences the weight allocated to the adviser, i.e. her estimated knowledge or reliability in disclosing an outcome.

In this study, we address these questions through a novel paradigm, inspired by previous work (Behrens et al., 2008; Cook et al., 2014) and designed to disentangle the relative importance of socially- and personally-acquired information in a decision. Socially-acquired information was operationalized in the form of advice on binary choices between options A and B. Advice was provided by two social peers (confederates), and were closely matched in fidelity, volatility, and content, but had different framings. One adviser often gave advice with an appetitive frame, such as “Choose A”. The other adviser often gave advice with an aversive frame, such as “Don’t choose B”. The task drives participants to track the history of reward for their choices, as well as the history of advisers’ fidelity, and integrate those two sources of information when making their choice. The framing of advice can affect participants’ estimation of each adviser’s fidelity, participants’ weighting of the two types of advice in their choice, or both. Computational modeling techniques, such as Bayesian and reinforcement learning modeling, enable us to see the world from the decision maker’s perspective and calculate the estimated trustworthiness of each adviser, as well as the expected reward value for each option. Combined with functional magnetic resonance imaging (fMRI), these models allowed us to identify brain regions tracking computational parameters required for performing this task (Daw and Doya, 2006; Cohen et al., 2017), and understand how framing of advice modulates those neuro-computational mechanisms. This enabled us to disentangle whether framing of advice influences estimation of knowledge of the two advisers, or their trustworthiness, or both, in regions involved in social learning and valuation of choice, such as TPJ and vmPFC, respectively.

## Results

### Experimental setup

Thirty participants (16 women, average age = 23.3 years) underwent fMRI while performing a decisionmaking task in which they chose between blue and green options to accumulate monetary rewards. On every trial, they also received an advice from one of two advisers about which option they should or should not choose. On every trial, the adviser could help the participant by revealing the correct option or could lie. Furthermore, the adviser framed their advice in either an appetitive manner (‘choose blue’) or an aversive manner (‘don’t choose green’) while keeping the content of the advice the same. While both “choose blue” and “don’t choose green” point to the same choice for the participant, they have different frames. The advisers were social peers (confederates, one marked in yellow, the other in purple) who participants met before the imaging session began, in a practice session. The practice session was designed to expose participants to the fact that both advisers would attempt to vary fidelity of their advice from trial to trial (i.e. to lie on some trials, but not systematically), in order to earn monetary rewards. However, unbeknownst to participants, during the experiment the advice from both confederates were generated using a computer program, with matched fidelity and volatility throughout the experiment. The difference between advice of the two advisers was only in the frame of their advice. One adviser gave appetitively framed advice in the majority of trials (appetitive adviser) and the other one mostly gave aversively framed advice (aversive adviser).

Participants differentiated between the two advisers, and that difference had behavioral consequences. During the imaging session, participants’ choices followed the appetitive adviser more frequently (t(29)=2.23, p=0.034; Figure 1c). After the imaging session, participants reported the trustworthiness of each adviser (explicitly, on a five-point scale), and their fairness (implicitly, through a variant of the Ultimatum Game (Sanfey et al., 2003; Tabibnia et al., 2008; Güroğlu et al., 2010)). Differential trustworthiness correlated with how closely participants’ choices followed the appetitive adviser more than the aversive one (Spearman correlation: r=0.54, p=0.002, Figure 1d), and there was also a strong correlation between perceived trustworthiness and fairness of the advisers (Supplementary Figure 1).

**Figure 1.**
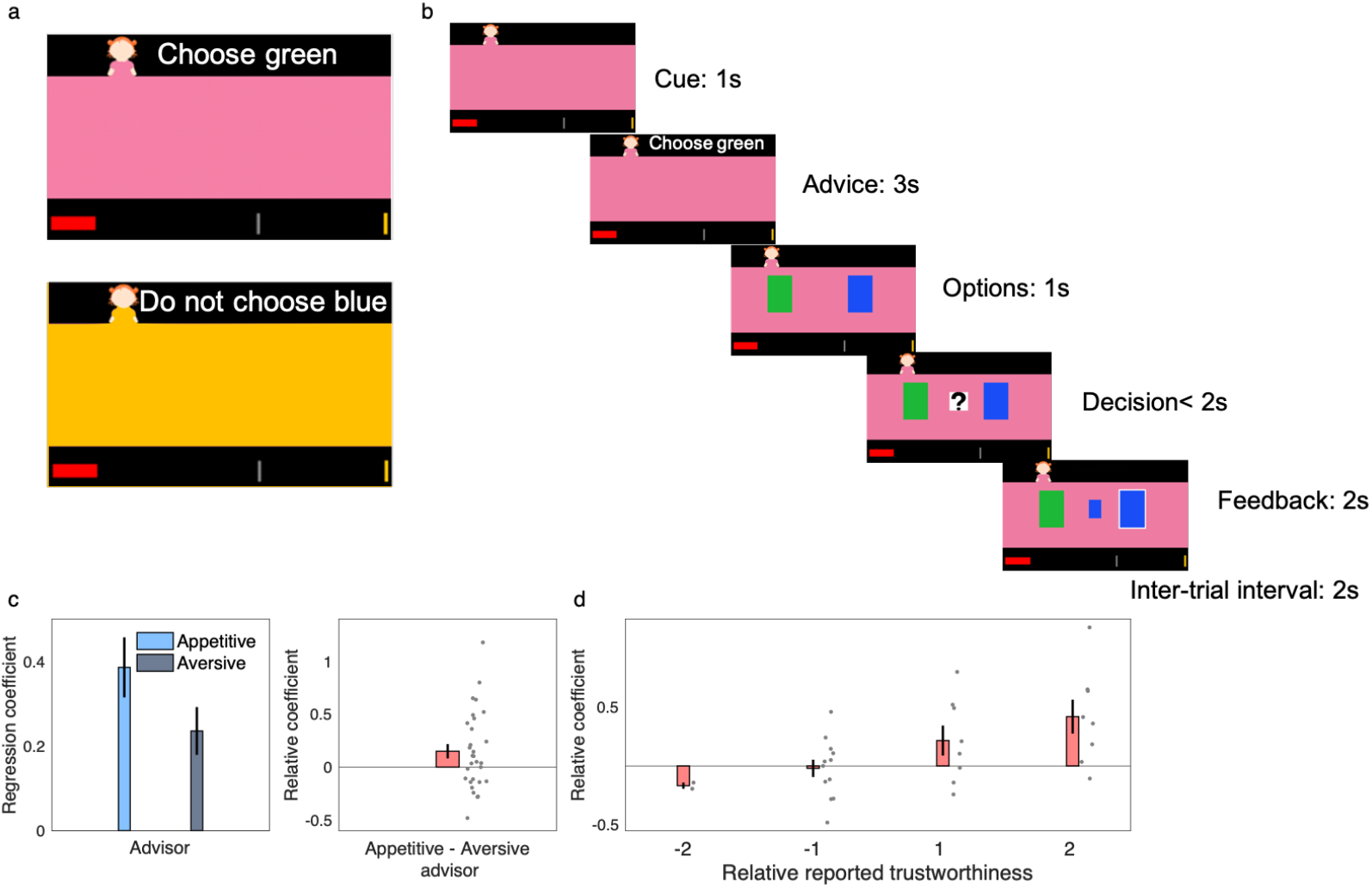
Experimental paradigm and behavioral finding. a) Participants received advice from one of two advisers cued with different colors. One adviser often gave appetitively framed advice, such as “Choose green” and the one often gave aversively framed advice, such as “Do not choose blue”. b) Timeline of the experimental task. On every trial, participants are presented with a color cue and then receive a written advice from the adviser. After choosing (button press) between two decision options, the correct choice (feedback) is revealed, and the total score (red bar at the bottom) increases with each correct response. c) Across all trials, participants followed the appetitive adviser more often than the aversive one, with a correspondingly positive average in the inter-participants distribution of the relative coefficient defined as the regression coefficient of appetitive-minus that of aversive-adviser. d) At the end of experiment, subjects were asked to rate trustworthiness of each adviser on a five-point scale. Across participants, relative reported trustworthiness was positively correlated with the relative coefficients of following advisers during the experimental task. Error bars are standard errors of the mean.

### Computational elements of choice under advice

Modeling techniques identified the computational mechanisms underlying the choice data. Optimal behavior in this task requires participants to combine social information (i.e. the advice, given the history of advisers’ fidelity) with the history of correct options (i.e. which color has been more rewarding recently). Both sources of information were present on every trial outcome, but their statistics were independently manipulated throughout the experiment (Figure 3ab). Importantly, in addition to fidelity, volatility of the social information was closely matched for the two advisers. Therefore, optimal decisionmaking requires the decision maker to track i) the probability that the advice is correct, ii) the probability of one color being rewarded throughout the experiment, and iii) to combine the two probabilities regardless of the current adviser. We considered a Bayesian learning model, the volatile Kalman filter (VKF), an extension of the Kalman filter to volatile environments such as our task (Piray and Daw, 2020). This model assumes that the first- and second-order statistics of observations (e.g. fidelity and volatility of advice) change over time. Similar Bayesian models have been used to model choice data in probabilistic paradigms similar to the current task (Behrens et al., 2008; Diaconescu et al., 2014). The VKF predicts the probability of each option being correct given the sequence of previous outcomes, and the probability that the advice is correct (Figure 3ab). These two sources of information were then employed to generate the probability of each option by combining prior belief parameters and the predicted probabilities (likelihood) according to Bayes rule. We assumed that different belief *weight* parameters encode subject-specific beliefs about the appetitive and aversive advisers, respectively (*w_ap_, w_aν_*). Higher values of the weight parameter were associated with higher probability of using social information for the trials associated with the corresponding adviser. Probability of using reward information was always the inverse of using the social information.

**Figure 2.**
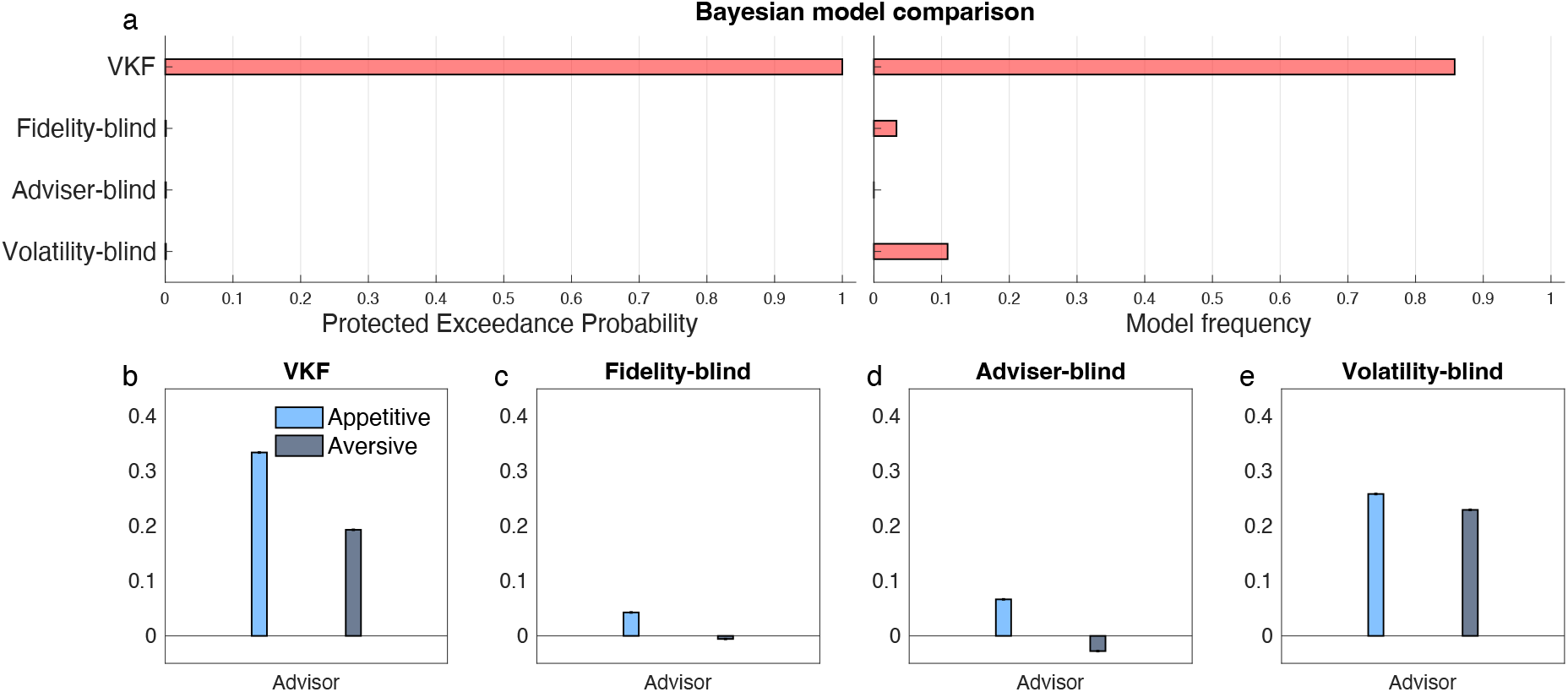
Model comparison results. a) Bayesian model comparison results are reported. The VKF model with different weights for the advisers was better than three simpler models: an advisers’ fidelity-blind model in which subjects followed the advice (possibly with different weights for each adviser) regardless of their fidelity history; an advisers’ frame-blind model in which subjects blindly followed the advice, possibly with different weights for each frame; and an adviser’s volatility-blind model with a simpler learning strategy (constant learning rate) with different weights for the two advisers. Protected exceedance probability (probability that the model being the most frequent at the group level taking into account the null possibility that none of the models account for data) and model frequency (ratio of participants explained by each model) are reported. b-e) Model simulation results for all models are reported, in which artificial datasets were created by the model (given the mean fitted parameters across subjects). Whereas the VKF model reproduce an effect similar to those seen in empirical data (i.e. Figure 1c), other models fail to reproduce such effects or they are worse than the VKF. In b-e, 5000 artificial datasets were generated per model. These datasets were then subject to the same logistic regression analysis as reported in Figure 1c. Mean and standard error of the mean are plotted in b-e. Errorbars might be too small to be visible.

**Figure 3.**
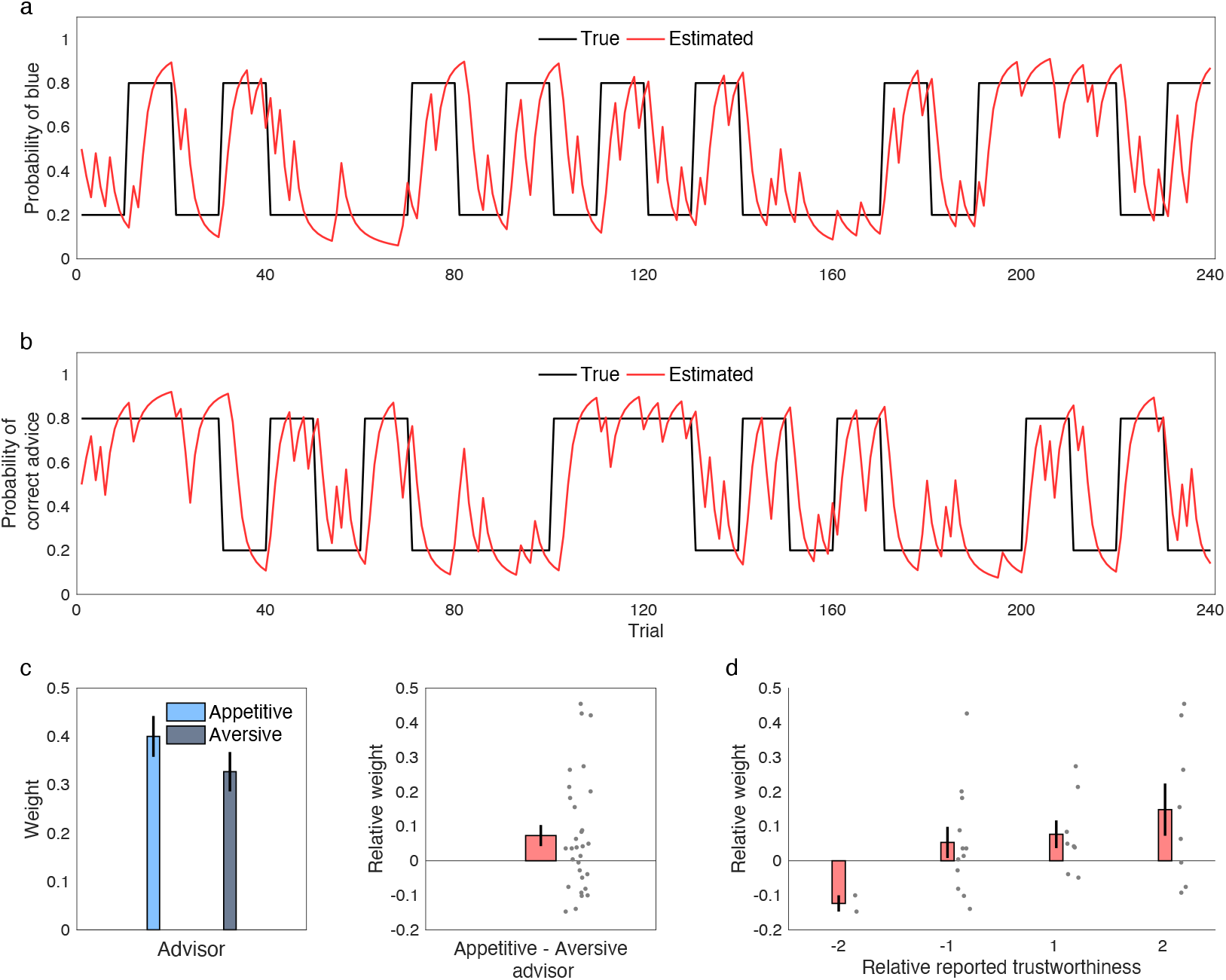
Computational findings. a, b) Probability schedules for the reward- and advice-based information. The black line shows the true probability of blue being correct (a) and the true probability of the advice being correct (b). The red line shows the VKF model’s estimate of the probabilities. c) Weight parameters from the winning model, in which weights represent the degree that subjects employed social information on trials associated with the appetitive and aversive adviser. Participants significantly relied more on the appetitive than the aversive adviser. The distribution of the relative weight parameters (appetitive minus aversive) is also shown. d) Relative weight as a function of relative reported trustworthiness is plotted. Error bars are standard errors of the mean. See also Supplementary Figures 3-4.

The VKF model provided a better account of the choice data than a number of alternative models. We compared the VKF model with alternative simpler models (Figure 3c) using random effects Bayesian model comparison tools (Piray et al., 2019a). First, the VKF model was better than a simpler strategy, in which subjects followed the advisers blindly with different prior belief parameters, but regardless of the history of advisers’ fidelity (Figure 2a – Fidelity-blind model). Second, the VKF model was better than another cognitively-lean strategy, in which subjects followed the advice, possibly with different weights for appetitive and aversive frame, but regardless of the adviser (Figure 2a – Adviser-blind model). Finally, the VKF model was better than an alternative learning model, in which the learning strategy was simpler by tracking only the first order statistics regardless of volatility (Figure 2a –Volatility-blind model). Statistics of fitted parameters of the winning model are reported in Supplementary Table 1. We then conducted a simulation analysis (Figure 2b-e) to study how well these models reproduce the behavioral effects seen in the empirical data (i.e. Figure 1c). Thus, 5000 artificial datasets were created in which choice was generated based on the models (using the fitted parameters). The VKF model was able to reproduce the main effect of interest (i.e. Figure 1c), but the blind models failed to capture those effects.

We then focused on parameters of the VKF model. First, we conducted a recovery analysis and confirmed that the VKF model parameters were recoverable (Supplementary Figure 2). Next, we studied the belief parameters of the VKF. The fitted belief parameters indicated that participants’ choices were more strongly influenced by the appetitive adviser. Individual values of the weight parameters, *w_ap_* and *w_aν_*, indicated that participants, on average, relied less on aversive advisers: the weight parameter for using social information was significantly higher for the appetitive adviser than for the aversive one (t(29)=2.40, P=0.023; Figure 3c). Furthermore, whereas the weight probability of using reward history for making decisions for the appetitive adviser was not significantly different from 0.5 (t(29)=2.02, P=0.053), this probability was strongly higher than 0.5 for the aversive adviser (t(29)=3.83, P<0.001). This observation indicates that participants used their own experience more than the social information when the advice was given by the aversive adviser. Moreover, individual differences in the weight parameters predicted reported trustworthiness of the two advisers. Specifically, the difference between the two belief weight parameters was correlated with the degree to which the appetitive adviser was reported to be more trustworthy than the aversive one (Spearman rank correlation: r=0.36, P=0.049; Figure 3d).

We conducted further analyses to investigate whether the different influence of the two advisers on participants’ choices was a consequence of differences in learning the advisers’ fidelity. We tested whether the association between the aversive advice (‘do not choose green’) and the recommended action (i.e. the blue one) might be harder to learn than the association between appetitive advice (‘choose blue’) and the recommended action (i.e. the blue one). First, we performed a logistic regression analysis, similar to the analysis presented in Figure 1, with two additional regressors evaluating the impact of fidelity on the previous trial and its interaction with the identity of the adviser. This analysis revealed a highly significant effect (t(29)=7.43, P<0.0001) of fidelity on the previous trial on current choice (i.e. one-trial social learning) indicating that participants processed and used advisers’ fidelity in their choice, but no significant interaction between this learning regressor and the identity of the adviser (t(29)=–1.63, P=0.11), indicating no difference in social learning between the two advisers (Supplementary Figure 5). Second, a model-based analysis confirmed and extended these one-trial learning effects by using full models and Bayesian statistics. Specifically, two additional models were fitted to choice data to investigate whether subjects processed the history of the two advisers differently. In particular, these extended VKF models considered whether different amount of noise governed learning processes for the two advisers. Note that VKF contains two different noise parameters that encode different types of noise. When compared with the original model in which the same noise parameters govern learning for both advisers, the extended VKF models would capture the presence of any differences in the learning process between the two advisers (e.g. the possibility that participants viewed the aversive adviser as more noisy). Bayesian model comparison analysis revealed that the original VKF model with no additional parameter accounts better for choice data (model frequency = 0.94; protected exceedance probability = 1), confirming that participants did not process the history of the two advisers differently (Supplementary Figure 5).

All computational analyses presented above assumed that the weight parameters for reward and social information sum to 1 (e.g. the weight for reward information on trials with the aversive adviser is 1 – *w_aν_*). Thus, the observed smaller weight for using the social information on trials with the aversive adviser inevitably means larger weight for using reward information on those trials. This suggests that there is an interaction effect between using social vs. reward information and the appetitive vs. aversive adviser. Therefore, we performed further analyses to test this interaction effect statistically. We considered a new VKF model identical to the original one but with four weight parameters (i.e. assuming different weight parameters for the reward and social information.) We then compared this model with the original model (we also considered a third mixing strategy in which the original VKF was extended to contain two different bias parameters for following the two advisers blindly; see Supplementary Information) using the same Bayesian model comparison procedure as above. This analysis revealed that the original model accounts better for choice data (Figure 4a), which confirms the original conclusion that the interaction was statistically significant (because the original model that hard-coded the interaction defeated the new model.) Nevertheless, we also analyzed the fitted weight parameters of the new model (Figure 4b-d). Consistent with the model comparison results, this analysis also revealed a significant interaction between using social vs. reward information and the appetitive vs. aversive adviser (t(29)=2.11, P=0.044). Post-hoc tests suggested that this effect is mainly driven by different weights for using the reward information for appetitive and aversive advisers (t(29)=–2.25, P=0.033), although there was also a trend for different weight for social information (t(29)=1.71, P=0.097). As anticipated, the interaction effect in the new model is strongly correlated with the interaction effect (i.e. double *w_ap_* – *w_aν_*) in the original model.

**Figure 4.**
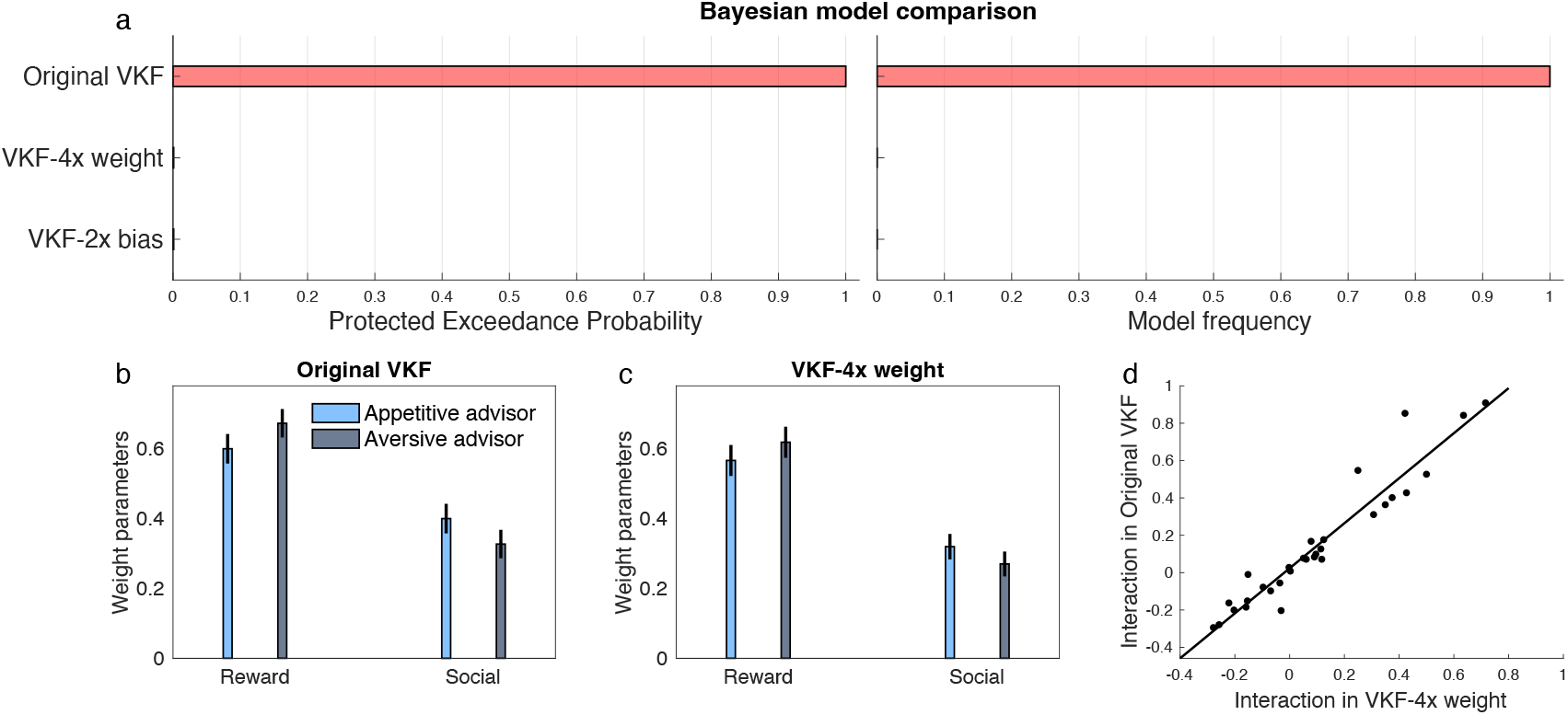
a) The original VKF model was compared with a new VKF model with four different weight parameters (VKF-4x weight), which allows the interaction effect between source of information (reward vs social) and adviser (appetitive vs aversive) to be tested. We also considered a third model that extends the original VKF by assigning different bias parameters for the two advisers (VKF-2x bias). Model comparison metrics showed that the original VKF is a more parsimonious account for the choice data. Both protected exceedance probability as well as model frequency of the original VKF were indistinguishable from 1. b-c) Weight parameters for the original model and the new VKF-4x model are plotted (note that for the original model, reward weights are defined as the inverse code of the social weights). The weights parameters of the VKF-4x model show a significant interaction between source of information and the adviser. c) scatterplot of interaction effect in the original model and the new VKF-4x model. Mean and standard error of the mean were plotted in b-c.

These computational analyses indicate that participants combined social advice and personal reward experience, relying more on the reward information and less on the social information on trials in which the advice was given by the aversive adviser. This effect can theoretically be mediated by neural processes at the time of choice, or by neural processes at the time of receiving feedback, or a combination of both. Using fMRI, we isolated brain activity associated with computational parameters quantifying reliance on social and reward information, during choice and as a function of learning.

### Choice-related effects in vmPFC

We focused on a ventromedial portion of the prefrontal cortex (vmPFC) that has been shown to combine social- and reward-related information at the time of choice in value-based decision-making (Behrens et al., 2008). We considered two decision-related regressors, time-locked to the presentation of the choice options. One regressor was based on social advice (social decision variable, SDV), and it was equal to 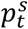 (estimated probability of advice being correct on trial *t*) and 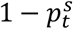 on trials in which the subject chose to follow and not follow the advice, respectively (see Methods – Computational model). The other regressor was based on personal experience of reward (reward-based decision variable, RDV), and it was equal to 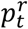 (estimated probability of blue being correct on trial *t*) and 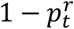 on trials in which the choice was the blue and the green option, respectively. Activity of the vmPFC was significantly correlated with a combination of both regressors (Behrens et al., 2008), consistent with its role in value-based decision making (Figure 5a; peak at x=−6, y=36, z=−18; voxel-level familywise small-volume corrected, k=713, t(29)=5.73, P_FWE-SVC_<0.001; see also Supplementary Figure 6). Post-hoc tests revealed a stronger association with the reward-based information (RDV) when the aversive adviser was involved (Figure 5b; t(29)=2.45, P=0.021), but no significant difference on socially-acquired information (SDV) (t(29)=0.52, P=0.52). Furthermore, there was inter-participant variation in the extent to which the vmPFC signal reflected personally-acquired information when the aversive or the appetitive adviser was involved. Crucially, variation in vmPFC signal predicted variation in the weight parameter capturing reliance on reward-based probability in the computational analysis (Figure 5C; r=0.39, P=0.035). Overall, these results suggest that participants relied more on their own experience on trials involving the aversive adviser.

**Figure 5.**
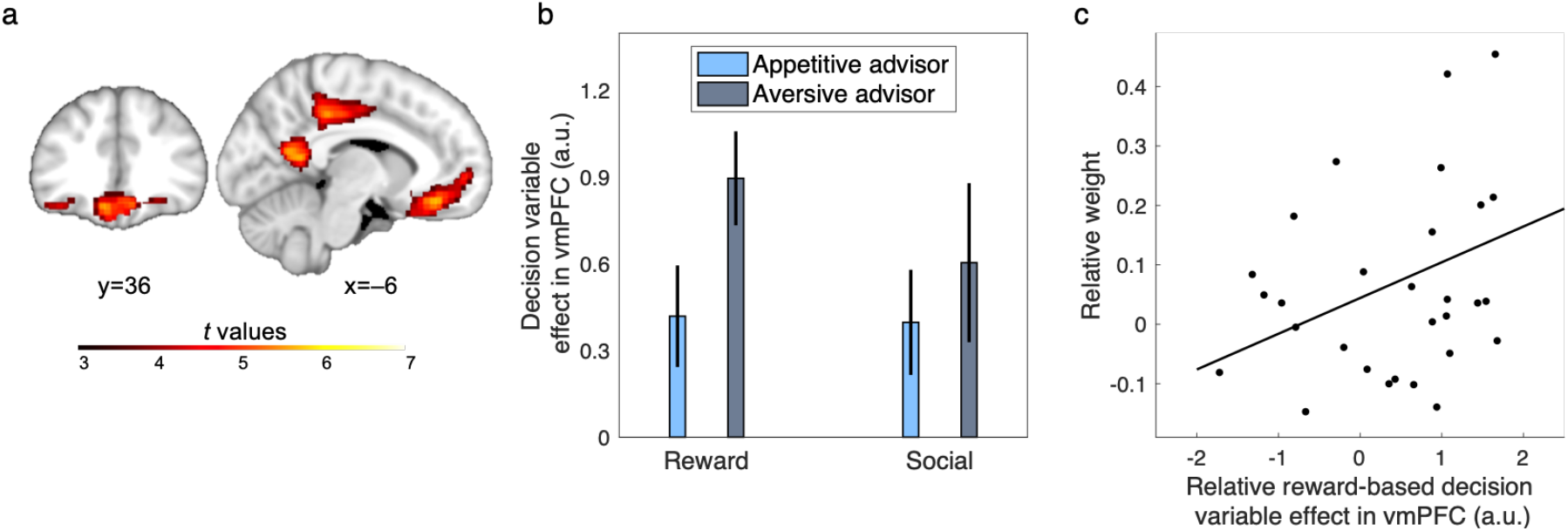
Decision-related activity in the vmPFC. A) Activation for the combination (mean contrast) of RDV and SDV at the time of decision making, when options were presented (thresholded at *P*<0.001 for illustration purposes). There is a significant effect in the vmPFC. B) Beta coefficients across all voxels of the VMPFC with uncorrected *P*<0.001. Decision-related activity of the vmPFC is significantly stronger for reward-based decision variable on trials associated with the aversive adviser. C) Inter-individual variability in behavior (differential use of reward-based information between appetitive and aversive advisers) was associated with neural variability in vmPFC (differential signal of personally-acquired information between appetitive and aversive advisers, as in panel B). Errorbars are standard errors of the mean. Unthresholded statistical maps are available at https://neurovault.org/collections/LINWGQZZ/. See also Supplementary Figure 4.

### Outcome-related effects in TPJ

We focused on a caudal portion of the TPJ that has been shown to track social prediction error during learning (Saxe, 2006; Behrens et al., 2008; Carter et al., 2012; Diaconescu et al., 2014). We considered two learning-related regressors, time-locked to the presentation of the outcomes. One regressor was based on the social prediction error (SPE), and it was given by the difference between observed and expected fidelity of the advice, weighted by the corresponding estimated uncertainty. The other regressor was based on the reward-based prediction error (RPE), which was given by the difference between observed and expected reward, weighted by corresponding estimated uncertainty. This analysis revealed that the right TPJ was significantly correlated with the social prediction error (Figure 6a; peak at x=42, y=–58, z=28, voxel-level familywise small-volume corrected, k=27, t(29)=4.94, P_FWE-SVC_=0.014). This effect was not significantly different for the appetitive and aversive advisers (across all voxels in the TPJ mask: t(29)=– 1.25, p=0.22). These results concur with the behavioral and computational analyses, confirming that the TPJ, a brain region known to track social prediction error, followed the fidelity of both advisers equally well. In other words, these results suggest that observed behavioral differences in the weight given to the two advisers are not driven by differences in ability to track their fidelity. In fact, if anything, the right TPJ was more active for the aversive adviser. There was also a significant effect of reward prediction error in TPJ, but only in the left hemisphere (Figure 6b; peak at x=–64, y=–32, z=44, voxel-level familywise small-volume corrected, k=96, t(29)=5.06, P_FWE-SVC_=0.006). (see also Supplementary Figures 7-9 for RPE effects in other areas of the brain, as well as volatility effects).

**Figure 6.**
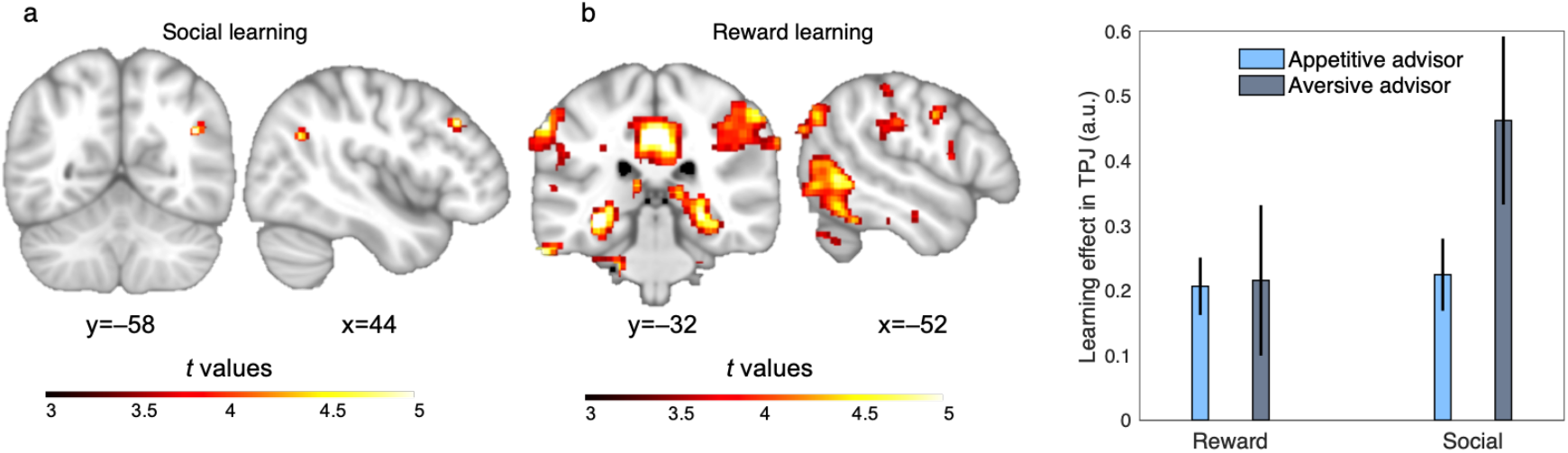
Learning activity in the TPJ. a) Activation for the SPE at the time of feedback, when it was revealed whether the advice was a lie or not. There is a significant effect in the TPJ. b) Activation for the RPE at the time of feedback. There is a significant effect in the TPJ, although this effect is in an anatomically distinct area compared with the social learning effect. Brain images are thresholded at *P*<0.001 uncorrected for illustration purposes. There was no significant difference between the two advisers regarding these effects. c) Beta coefficients across all voxels of the TPJ mask with uncorrected *P*<0.001 for both types of learning are plotted. Unthresholded statistical maps are available at https://neurovault.org/collections/LINWGQZZ/.

## Discussion

Humans operate in a hyper-cooperative social environment, where individuals offer their knowledge to non-kins in the form of advice (Boyd and Richerson, 2009; Salali et al., 2016). Accordingly, we need computational and neural mechanisms to adaptively combine personal experience with socially mediated knowledge. Here, we identify neuro-computational characteristics of a bias in how pieces of advice are combined with personally-acquired knowledge. The bias leads participants to rely less on aversively-than appetitively-framed advice to make decisions, despite matched fidelity, volatility, and content of the advice. This social framing bias occurred at the time of choice, with a correspondingly stronger weight allocated to personal experience over aversively-framed advice when those sources of information are combined in the vmPFC. These findings suggest that social advice is systematically incorporated into the decision-making process. Fittingly, this effect was found in the vmPFC, a region known to combine volatile knowledge about social partners with personal knowledge (Stolk et al., 2015), and to guide current actions by updating the relative value of different models of the world (Nicolle et al., 2012).

Several studies in psychology, neuroscience and behavioral economics have shown that human decision making is influenced by how the decision problem is framed (Kahneman and Tversky, 1979; Kahneman et al., 1982; De Martino et al., 2006; Rangel et al., 2008). The framing effect has been often used as an example of how emotional valence might play a disruptive role in rational decision making. In that perspective, it could be argued that the current findings are a straightforward generalization of well-known Pavlovian biases in individual value-based learning (Guitart-Masip et al., 2012) to socially-mediated learning. However, that interpretation does not account for the observation that the social framing bias led participants to trust and perceive as fairer those advisers that provided predominantly appetitively-framed advice, without biasing participants’ confidence in the knowledge of the advisers, and without down-weighting their influence. An alternative viewpoint suggests that socially-driven modulations of decision making might stem from appraising how an advice is communicatively embedded (McKenzie and Nelson, 2003; Sher and McKenzie, 2006; Frith and Singer, 2008). Namely, considering the communicative expectations generated by the advice might offer a more complex but comprehensive interpretation of the findings, accounting for social framing bias, matched confidence in advisers’ knowledge, yet different trust in them. The two types of advice, while informationally matched, impose different cognitive demands on the participants. The aversively-framed advice imposes on the participant the need to resolve the negation and infer the suggested choice. Imposing this marginal but un-necessary cognitive demand on the recipient of the advice violates the cooperative principle of human communication (Grice, 1975), and violates the expectation of additional pragmatic implications generated in the listener when that communicative principle is ignored.

The social framing bias found in the current work adds to a line of research in social psychology and neuroscience, which shows that people exploit social information less than optimal. In fact, people exhibit different types of biases in using social information, such as equality bias (i.e. the tendency to allow everyone an equal say in collective decisions) (Mahmoodi et al., 2015; Hertz et al., 2016), egocentric bias (i.e. the tendency to rely more on their own individual information) (Yaniv and Kleinberger, 2000; Heyes, 2016), confirmation bias (i.e. the tendency to discount social information that undermine their past judgements) (Kappes et al., 2020), and reciprocity bias (weighting others’ opinion by the weight they give to our opinion) (Mahmoodi et al., 2018). Optimal decision making in this task requires equal weight for both advisers regardless of the frame of advice as the content of information was matched for the two advisers. In fact, to ensure that our results are not contaminated with differences in content of advice, we matched both fidelity and volatility of the two advisers in small bins of 10 trials.

But why do people show such a bias? An important aspect of our experimental paradigm was the fact that participant made binary choices. This was important because such a binary choice guaranteed that the content of information was matched for both frames (i.e. that “Choose green” is equivalent to “Do not choose blue”). However, real-life decisions often require choosing between more than two options, and in such environments, an aversively framed advice is less informative for decision making than an appetitive frame. In naturalistic settings, it could be argued that an appetitively-framed advice, on average, leads to faster and better decisions, possibly resulting in a systematic bias even when choosing between equivalent options. On the other hand, aversive frames are more likely to be consequential for survival, i.e. to signal threats, in particular when combined with nonverbal bodily signals. Our data are clearly consistent with the former possibility. It remains to be seen whether threat-related social signals might alter the social framing effect reported here, e.g. by influencing computations at the time of learning rather than choice, as observed in reinforcement learning studies (Piray et al., 2019b).

Our results corroborate the separate contributions of different social brain systems in tracking and integrating advice: whereas right TPJ was engaged in social learning by computing a social prediction error, vmPFC contributed in integrating the social information with the personally-acquired feedback-based information. This distinction is consistent with previous work on neural mechanisms underlying strategic influence in advice giving (Hertz et al., 2017), in which the right TPJ was associated with an internal social comparison process, and medial PFC was associated with evaluation of social outcomes in relation to the self. A similar distinction has been found in two-person games, in which the right TPJ was associated with computations regarding the behavior of others, and the medial PFC was associated with computing the internal valuation process given the degree of social influence expected(Hampton et al., 2008). However, based on our findings, the role of vmPFC could equally well be interpreted as a general neural mechanism for valuation, regardless of the fact that one source of value was social advice here. This interpretation is consistent with the neuroeconomics and decision neuroscience literature (Hampton et al., 2007; Kable and Glimcher, 2007; Behrens et al., 2009; Haber and Knutson, 2009, 2010; Hare et al., 2009), in which vmPFC is shown to encode a common valuation signal.

Our results suggest that human participants are adept in tracking mental state of their social peers even when their behavior is volatile. Statistical principles indicate that for optimal learning, organisms should dissociate volatility from a second type of noise, i.e. moment-to-moment stochasticity (Piray and Daw, 2021). These theories also show that these two types of noise should have opposite effects on learning: whereas volatility should increase the learning rate, stochasticity should decrease the learning rate. While effects of volatility on social learning has been well studied (Behrens et al., 2008, 2009; Diaconescu et al., 2014, 2020; Reiter et al., 2017), there is currently no work, to our knowledge, that has studied dissociation between volatility and stochasticity in social learning. Similarly, dissociating the two types of noise is crucial for advice taking: while volatility means that speed of changes in a social peer’s mental state, stochasticity means that social peer’s advices are noisy but not necessarily changing in any meaningful way. Importantly, these factors are interdependent as they are alternative explanations for experience noise, which makes the problem computationally more challenging. Moreover, framing of advice might have influence these processes separately. It is remained for future studies to expand the current design to study stochasticity and its interactions with volatility in social learning.

## Methods

### Experimental paradigm

The experimental paradigm and the cover story were taken from (Behrens et al., 2008). In this study, verbal advice was given to subjects by two confederates: one confederate often used an appetitive frame to give advice (in 80% of trials) and the other confederate often used an aversive frame to give advice.

Participants performed a probabilistic decision-making task in the scanner, in which they chose between blue and green tokens to accumulate reward (i.e. 1 or 0 point). Participants were instructed that on every trial, one of the colors leads to reward, while the other one leads to no reward. Therefore, it was clear for the participants that the probability of win given blue and green was inverse of each other, i.e. *p*(win|blue) = 1 – *p*(win|green). Participants’ accumulated rewards were shown on the screen using a red score-bar, the length of which was proportional to the accumulated reward. A gold and a silver target were also shown on the screen and subjects were instructed that if they land their red score-bar in either of those targets, they receive 8 and 4 euros bonus, respectively. On each trial, the subject was first presented with a color cue (yellow or purple) and a cartoon cue indicating the adviser of this trial. Participants were instructed that each color is associated with one adviser throughout the experiment. One second after seeing the cue, the subject received a written advice (in Dutch), which could take 1 of 4 forms: “Choose blue”, “Choose green”, “Don’t choose green” and “Don’t choose blue”. The advice remained on the screen for 3 seconds to ensure that subjects have sufficient time to process the advice. Next, two options (blue and green tokens) appeared on the screen for 1 second, during which subjects were not allowed to make their choice. This was followed by a question mark on the screen, indicating that subjects were allowed to make their choice (in less than 2 seconds). The correct option on that trial was then revealed on the screen .25 second after subjects made their choice and remained onscreen for 1.75 second. If the subject’s choice was correct, a coin, and otherwise, a red cross appeared on the screen 1 second after the correct option was revealed. Trials were terminated by an inter-trial interval of 1.5-2.5 seconds (except three trials throughout the task with longer intervals of about 20 seconds).

### Cover story

The cover story was similar to that of (Behrens et al., 2008). Participants were introduced to two actresses before the experiment began and all three of them were brought to a behavioral lab in which they received instructions and practiced the task together. In the behavioral lab, subjects and confederates were instructed that there were two different roles in the experiment: adviser and receiver. Participants were always assigned as the receiver and confederates as advisers (using a counterfeit test). The advisers’ task was to give trial-by-trial advice and their goal was to land the receiver’s score bar in either a golden or a silver range to win 8 or 4 euro of bonus, respectively. Importantly, instructions about the advisers’ task were given in front of participants. Critically, advisers were told that they need to mix helpful and harmful advice to maximize their bonus reward.

With examples, it was made clear that advisers’ motive to give a helpful or harmful advice might change during the task given the receiver’s score-bar location and the length and location of the golden and silver ranges. For example, if the receiver’s score is far behind the golden range, the advisers are motivated to give her helpful advice to help her to reach to the golden range. However, as soon as the receiver is already in the golden range, advisers are motivated to give harmful advice to keep her in that range. The advisers were told that each trial will be randomly assigned to one of them, in which they should choose between four options as their advice: “Choose [correct]”, “Not choose [incorrect], “Choose [incorrect]” and “Not choose [correct]”. The advisers were also told that they don’t know which color is correct or incorrect, but their advice will be translated according to the correct color on that specific trial. For example, if they select “Choose [correct]” and the blue is the correct color on that trial, their advice which appears on the receiver’s screen would be “Choose blue”. Therefore, it was made clear that advisers can choose not only the content of their advice (correct or incorrect), but also the frame of their advice (choose or not choose). They were told that they will perform the experiment in different labs and they don’t see each other afterwards.

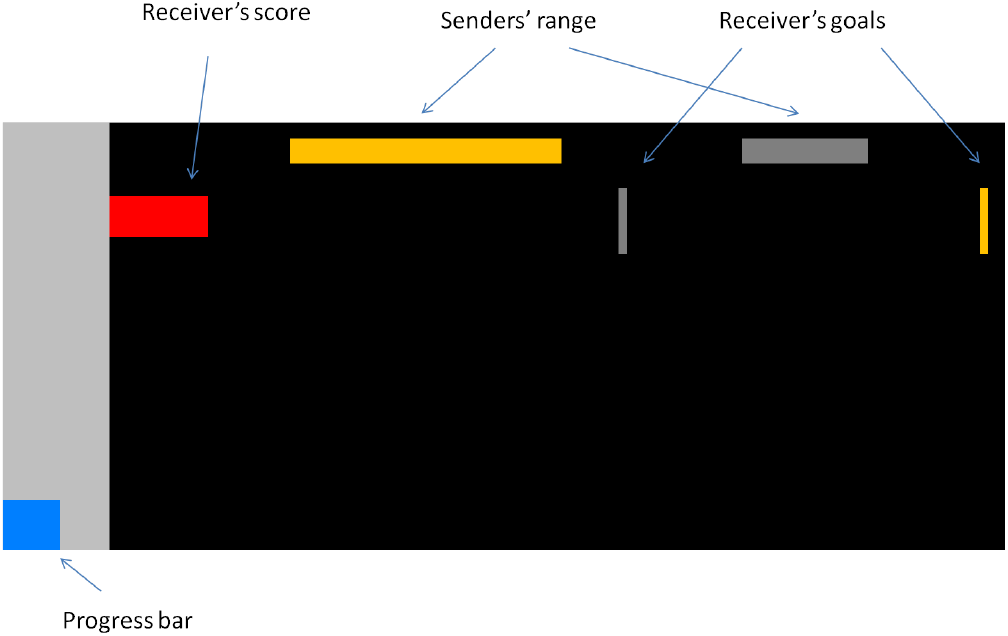
Advisers’ task. The two advisers were instructed and practiced their task (in front of the subject). They were told that their task was to use their advice to facilitate that the receiver’s red score-bar would land within their own golden range or a silver range, associated with 8 or 4 euros bonus. A progress bar signaled the number of trials passed. The advisers were told that their silver and golden ranges were the same and each trial would be randomly assigned to one of them.

The goal of the cover story was to make clear for the subject that 1) advisers have rational motives to give helpful or harmful advice in different phases of experiments (depending on the location and length of the golden and silver ranges); and 2) those motivations are the same for both advisers; 3) however, advisers’ advice is independent of each other (as they don’t see or communicate with each other). 4) Advisers’ advice is also independent of the subject’s choices, and, therefore, the subject can go against the advice or follow it as long as they want, 5) the advice is an extra and independent piece of information, because it is independent of the actual color being correct. Several examples were given during the instructions to make these points explicitly clear.

At the end of the experiment, subjects were asked to report which adviser they found to be more trustworthy. Next, they were asked to rate each adviser trustworthiness on a scale from 1 to 5. These data were used for correlation analysis in Figures 1–2.

### Computational modeling

The main model of interest in this study is a hierarchical Bayesian learning model, called volatile Kalman filter (Piray and Daw, 2020), extends the celebrated Kalman filter model to volatile environments in which both the first- and second-order statistics of the environment evolve over time. In addition to the error-correcting rule of Kalman filter and other classical models (e.g. Rescorla-Wagner model) for learning observations, the VKF also learns volatility according to a second error-correcting rule. We used the binary version of this algorithm here as observations are binary. Specifically, if *o_t_* is the observation on trial *t*, the VKF tracks the mean, *m_t_*, of the hidden state of the environment using an uncertainty-weighted prediction error signal:

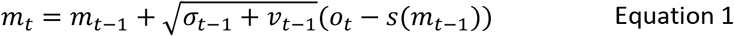

where *σ*_*t*–1_ is the estimated uncertainty on the previous trial, *ν*_*t*–1_ is the estimated volatility on the previous trial, *o_t_* is the current observation (binary: 0 or 1) and *s*(.) is the sigmoid function:

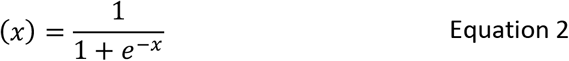

On every trial, VKF updates its uncertainty and volatility estimates:

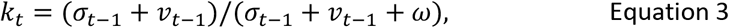

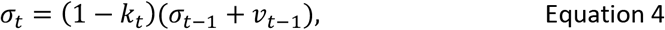

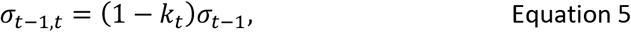

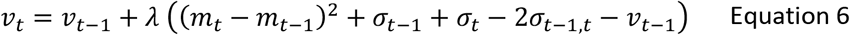

The VKF model requires three constant parameters: 0 < *λ* < 1 indicating volatility update rate, *ν*_0_ > 0 determining the initial value of volatility and *ω* > 0 indicating observation noise. Note that volatility estimates are updated using an (second-order) error correcting rule. Also, note that if *λ* = 0, then the VKF is equivalent to a (binary) Kalman filter in which the process variance parameter is *ν*_0_. Both mean and uncertainty of the VKF were initiated at 0.

For every subject, the VKF algorithm was used twice. First, we used VKF to quantify value of each option based on reward history, 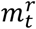. In this case, observations were the color of the correct option (1=blue, 0=green). Thus, larger values of 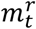 indicate subject’s estimated probability that the correct option is be the blue one. This is learnt using Equations 1–6 and based on three free parameters, *λ_r_*, *ω_r_* and *ν*_0_.

Second, we used the VKF to quantify value of each option based on the social experience and the advice, 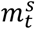. In this case, observations were the correctness of the advice (1=correct, 0=incorrect). Thus, larger values indicate subject’s belief that the advice should be followed. This is also learnt using Equations 1–6 and based on *λ_S_*, *ω_S_* and ν_0_.

Next, a choice model was used to generate probability of choosing each option based on 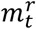 and 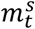. First, probability of the advice based on the social information was quantified using a sigmoid (softmax) function:

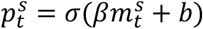

where *β* > 0 is the decision noise parameter and *b* is the value-independent bias parameter for following the advice. Second, we computed the probability of the advice given the personal information:

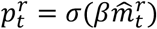

where 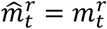 if the advice is the blue option (i.e. “Choose blue” or “Don’t choose green”) and 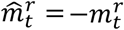 if the advice is the green option.

Finally, these two probabilities were combined to construct probability of following the advice using a mixture model with two prior belief parameters (weights):

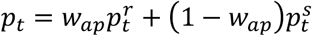 for the trials associated with the appetitive adviser
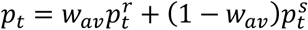 for the trials associated with the aversive adviser

where *w_ap_* and *w_aν_* indicate weight parameters corresponding to the trials associated with the appetitive and aversive adviser, respectively. Both *w_ap_* and *w_aν_* are constrained in the unit range and indicate the weight of using own experience. For example, if *w_ap_* = 1, the subject relies entirely on her own experience and ignores the social information. Conversely, if *w_ap_* = 0, the subject relies entirely on social information and ignores her own experience from past trials. In fact, this mixing strategy is reminiscent of mixture models in machine learning, in which one can assume that there is a latent binary variable on every trial, which is, say on appetitive trials, sampled based on probability *w_ap_*. If this variable is 1, the agent makes choice based on 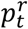, otherwise the choice is based on 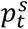. Across all trials (i.e. after marginalizing this latent variable), the probability of choice is given by above equations. This interpretation of the model based sampling a latent variable provides a computational rationale for employing this mixing strategies for implementing choice based on two independent sources of information.

Therefore, in total, this model has 9 free parameters: *w_ap_*, *w_aν_*, *β*, *b*, *ν*_0_, *ω*^S^, *λ^S^*, *λ^r^* and *ω^r^*. This is the base model that was referred as the VKF or original VKF in the main text.

In Figure 2, we considered three cognitively-lean models. First, we considered the possibility that subjects do not follow the intention of the confederate and follow advice (or go against advice) blindly, although with different weights for the advisers (fidelity-blind). For learning from reward feedback, this model is identical to original VKF. Therefore, this model has six parameters: *w_ap_*, *w_aν_*, *β*, *ν*_0_, *λ_r_* and *ω_r_*. We also considered a similar model in which different weights were given to appetitively- and aversively-framed advice but not the adviser (adviser-blind). Note that although this model has some overlap with the previous one, but is not identical to that one as advisers did not use their dominant frame in all trials. We also considered a simpler reinforcement learning strategy, which is based on only following the first-order statistics and fewer free parameters (volatility-blind). Instead of equations 1–6, the reinforcement learning model learns value of two actions, *a*_1_ and *a*_2_ (e.g. blue and green) by updating the value of chosen action using prediction errors:

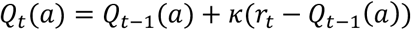

where *a* is the chosen action on trial *t*, *r_t_* is the reward on this trial, *Q_t_* is the action-value on trial *t* and *κ* is the learning rate (a constant parameter bounded in the unit range). We then define *m_t_* = *Q_t_*(*a* = 1) – *Q_t_*(*a* = 2). We used this algorithm twice: first with reward-related actions (*a* = 1 for choosing blue and *a* = 2 for choosing green) and *κ*_r_ to obtain 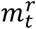; and then with advice-related actions (α= 1 for following the advice and *a*= 2 for not following) and *κ_S_*. to obtain 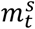. The same choice model as the VKF (containing a decision noise, a bias and two weight parameters) was then used to obtain probability of choices. Therefore, this model contains 6 free parameters: *w_ap_*, *w_aν_*, *β*, *b*, *κ_r_*, *κ_S_*.

We also performed an analysis for testing possible effects of advisers’ on learning over and above their direct effect on choice. This analysis confirmed the theory-neutral results (Supplementary Figure 5a) that subjects were able to follow both advisers equally well. For this analysis, we considered two models with additional parameters. These models were variants of the original VKF. The first model (VKF-2x *λ*), assumed that different volatility learning parameters (that also give the degree of noise at the volatility level) govern learning for the two advisers (*λ*_S_ was different for the two advisers). The second model assumed that different noise parameters, *ω_S_*, govern learning for the two advisers (VKF-2x *ω*). As reported in the Results and Supplementary Figure 5, Bayesian model comparison analysis revealed that the original model with no additional parameter provided a more parsimonious account of data.

Finally, two other models were considered with the same learning process as the VKF, but with different parametrization for mixing reward and social information. The first model had two additional weight parameters for the reward information. In particular, these two probabilities were combined to construct probability of following the advice four weight parameters:

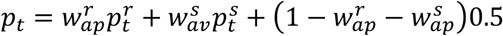 for the trials associated with the appetitive adviser
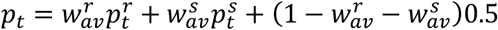 for the trials associated with the aversive adviser

Put it simply, these equations, for example for trials associated with the appetitive adviser, mean that with probability 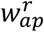 participants follow the reward information, with probability 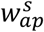 they follow the social information, and with probability 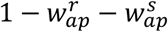 they choose randomly (i.e. probability 0.5 for each option). This is the VKF-4x weight model in Figure 4.

The second model was the same as the original VKF, with one different bias parameters for computing 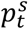 on trials associated with the appetitive vs aversive advisers. This is model VKF-2x bias in Figure 4.

### Model comparison and parameter estimation

We then used hierarchical Bayesian inference, HBI (Piray et al., 2019a), to compare these models. The HBI constructs subject-level priors based on the group-level statistics (empirical priors) and takes a random effects approach to model identity (different models might be the true model in different subjects). This method has been shown to be more robust than other methods for model selection (Piray et al., 2019a). This method fits parameters in the infinite real-space and transforms them to obtain *actual* parameters fed to the models. Therefore, appropriate transform functions were used for this purpose: the sigmoid function to transform parameters bounded in the unit range or with an upper bound and the exponential function to transform parameters bounded to positive value. For the VKF algorithm, we assumed an upper bound of 10 for the observation noise parameters (ω_+_and *ω_s_*) to have robust estimates. The HBI algorithm is available online at https://github.com/payampiray/cbm.

For subject-level analyses of the parameters, we used individually fitted parameters of model the original VKF. Unlike parameters estimated by the HBI that are regularized according to all subjects’ data, the individually fitted parameters are independently estimated and therefore can be used in regular statistical tests. These parameters were obtained using a maximum-a-posteriori procedure, in which the prior mean and variance for all parameters (before transformation) were assumed to be 0 and 3.5, respectively. This initial variance is chosen to ensure that the parameters could vary in a wide range with no substantial effect of prior. Specifically, for any parameter bounded in the unit range, the log-effect of this prior is less than one chance-level choice (i.e log0.5) for any value of that parameter between 0.1 and 0.9.

### fMRI acquisition and preprocessing

Functional images were acquired using a 3T Skyra MRI system (Siemens), using a multiband sequence(Moeller et al., 2010) with acceleration factor 3 (Flip angle = 60 degrees, FOV = 64×64×33 voxels, voxel resolution of 3.5 mm isotropic, TR/TE =1000/30 ms). Structural images were acquired using a T1-weighted MP-Rage sequence (Flip angle = 8 degrees, FOV = 192×256×256, voxel size 1×1×1, TR/TE = 2300/3.03ms). The task was presented and time-locked to fMRI data using Presentation (Neurobehavioural Systems, USA).

Data was preprocessed using FSL tools. Motion correction was applied using rigid body registration to the central volume and high pass temporal filtering was applied using a Gaussian-weighted running lines filter, with a 3dB cutoff of 100s. The T1-weighted images were spatially coregistered to functional images using linear (FLIRT) and non-linear tools (FNIRT). Noise effects in data were removed using FMRIB’s ICA-based Xnoiseifier tool(Salimi-Khorshidi et al., 2014), which uses independent component analysis (ICA) and classification techniques to identify noise components in data (using a trained version based on HCP_hp2000 dataset and R=10). We also measured physiological signals for the purpose of denoising fMRI data using RETROICOR (Glover et al., 2000). Pulse was recorded using a pulse oximeter transducer affixed to the middle or ring finger of the non-dominant hand and respiration was measured using a respiration belt. Recorded signals were then used to generate covariate nuisance regressors (25 in total) using a fifth-order Fourier model of the cardiac and respiratory phase-related modulation of the BOLD signal (Birn et al., 2006; Shmueli et al., 2007; van Buuren et al., 2009).

### fMRI analysis

We performed FMRI general linear model (GLM) analyses using SPM12. Similar to Behrens et al. (Behrens et al., 2008), we considered two GLM models, where the first looked for decision-related activity and the second for learning-related activity. In both GLMs, four sets of two regressors, each containing one regressor per cue type, were considered. These regressors were time-locked to the presentation of cues, advice, options and outcomes onscreen. In both GLMs, we also included 25 noise regressors obtained using the RETROICOR analysis of the pulse and respiration signals. Twelve motion regressors representing six motion parameters obtained from the brain-realignment procedure and their first derivative were also included.

In the decision-related GLM, two decision-variable parametric regressors, one based on reward information and the other one based on social information, were also included separately for each cue, both time-locked to presentation of options. The reward-based decision variable (RDV) was defined based on the interaction of 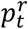 and the reward-related choice. Thus, RDV was equal to 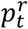 and 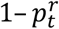 on trials in which the choice was the blue and the green option, respectively. The social decision variable (SDV) was defined based on the interaction of 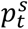 and the advice-related choice. Thus, SDV was equal to 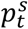 and 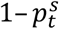 on trials in which the subject chose to follow and not follow the advice, respectively.

In the learning-related GLM, four parametric regressors were included, which were all time-locked to the presentation of outcomes. Two error regressors, an uncertainty-weighted reward-based prediction error (RPE) regressor (in the choice frame) and an uncertainty-weighted social prediction error (SPE) regressor were included (the update term in Equation 1). These regressors were defined based on equation 1. Two other learning regressors encoding volatility estimate were also included separately for each cue.

We used a common set of (group-level) parameters obtained by fitting a variant of M1 to data using the HBI to generate parametric regressors for the above GLM FMRI analyses. The only difference between this model and the original VKF model was the assumption that the weights are the same for both advisers. This is critical because it ensures that any difference in fMRI data is not simply due to having regressors generated based on different weights. Furthermore, any between-subject variability in coefficients of GLM FMRI cannot be attributed to parameters (correlation analysis of those coefficients and model parameters is valid and not circular). We used anatomical masks for region of interest (ROI) analyses according to recent connectivity-based parcellation studies(Mars et al., 2012; Neubert et al., 2015). Clusters 1 and 2 found by Mars et al. (Mars et al., 2012) was used as the TPJ mask (after smoothing with a 6mm kernel to avoid sharp edges). Cluster 4 of (Neubert et al., 2015) was used as the vmPFC mask.

