## Supplementary Materials for "Social framing effects in decision making"

### Supplementary results

Participants also performed a modified version of the ultimatum game after performing the main task. Participants were instructed that in this game, they play with one of the advisers on every trial, but they don't know which adviser is playing on each trial. They were also told that the adviser is tasked with splitting 1 euro between herself and the participant and if they accept the proposal, the money is split per the proposal (and both receive nothing if they reject). Critically, participants were asked to guess which adviser was playing on every trial. They were also instructed that they receive a monetary bonus for this game, which depends on the amount of money they accumulated on trials that their choice about adviser was correct. Unbeknownst to participants, however, all proposals were programmed.

We analyzed these data using two softmax models: one in which accept/reject choice data were only a function of the amount of the offer ($b_o$) (adviser-independent model); and one in which accept/reject choice was also a function of the trial-by-trial guesses about the adviser $b_a$ (adviser dependent model). We then used the same hierarchical Bayesian inference (HBI) techniques as we used for the other analyses to quantify evidence in favor of each model. The HBI regularizes individual fits according to the group statistics and, therefore, estimation of both models is tractable and finite for all participants. This analysis showed strong evidence in favor of the adviser dependent model, indicating that participants' guesses about the identity of the adviser was a predictive of the accept/reject choice data. As Figure S1 shows, subjects found the appetitive adviser fairer on average (positive $b_a$), although there were large individual differences in $b_a$. Importantly, however, individual values of $b_a$ were strongly correlated with the degree to which subjects reported the appetitive adviser to be more trustworthy than the aversive one at the end of experiment (r=0.78, p<0.000001). These data suggest that aversively framed advice develop trustworthiness biases about peers beyond the immediate context.

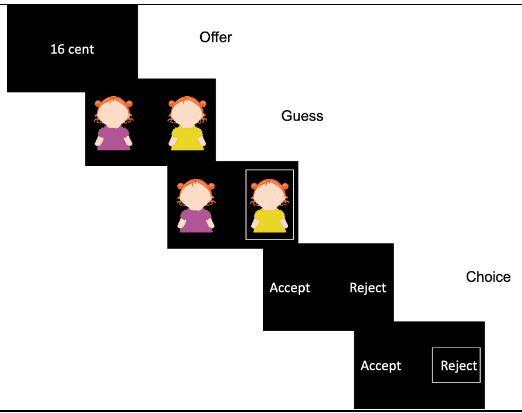


Supplementary Figure 1. A) participants performed a modified version of the ultimatum game. On every trial, participants saw an offer, made a guess about the adviser and then chose to accept or reject the offer. B) An adviser-independent model (AIM) and an adviser-dependent model (ADM), in which trial-by-trial guesses were also a predictor of the choice data, were

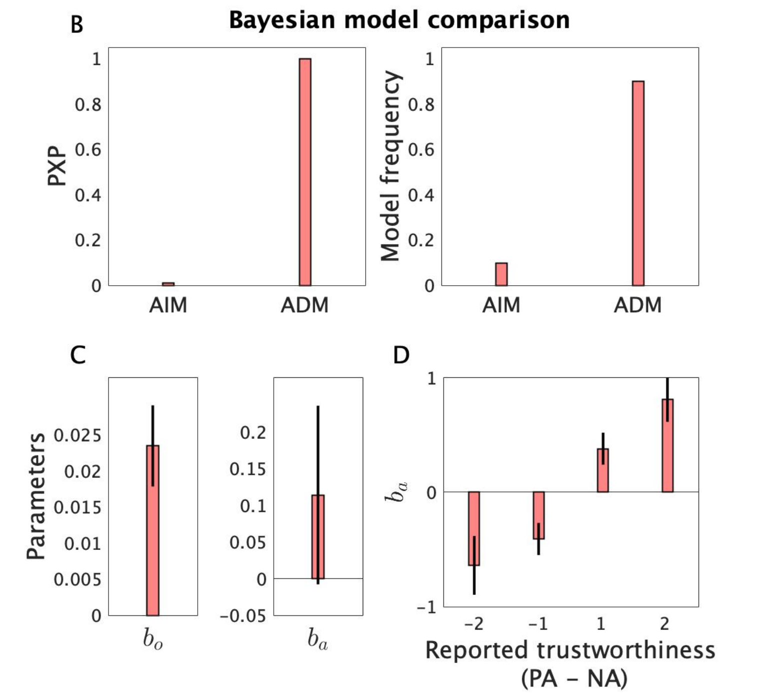


compared using Bayesian inference tools. Both protected exceedance probability and model frequency measures indicate that the ADM captures the data better than the AIM. Thus, participants' belief about the adviser is a predictor of their choice. C) On average, subjects showed positive $b_a$, which indicates a positive and negative tendency to accept the offer for the appetitive and aversive advisers, respectively. D) $b_a$ was correlated with the degree to which participants reported the appetitive adviser to be more trustworthy than the aversive one at the end of experiment.

#### Recovery analysis

We performed a recovery analysis for parameters of the winning VKF model presented in Figures 2-3. We simulated 100 artificial datasets (with the same number of trials as in the main task). We then applied the same computational fitting procedure used for fitting the empirical dataset to these artificial datasets. As shown in Supplementary Figure 2, parameters were quite well recoverable. In particular, the true parameter was always within the range of one standard deviation of the mean fitted parameter (the shaded area in Supplementary Figure 2). Importantly, recovery for the two critical weight parameters were almost perfect.

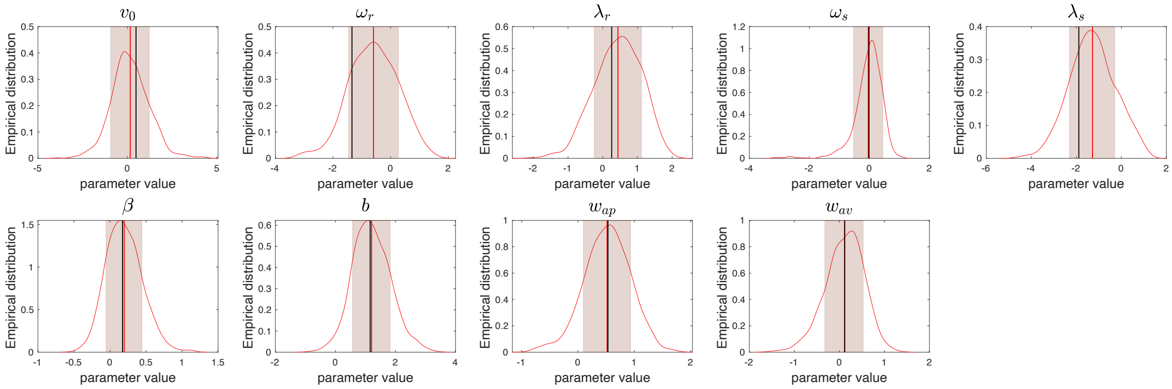


Supplementary Figure 2. Recovery analysis for the VKF model with two weight parameters. Empirical distribution for recovered parameters were plotted, along with the mean (red vertical line) and standard deviation (shaded area). The black vertical line is the true parameter, which always falls within the shaded area and often very close to the mean of empirical distribution. Importantly, recovery is almost perfect for the critical weight parameters ($w_{ap}$ and $w_{av}$).

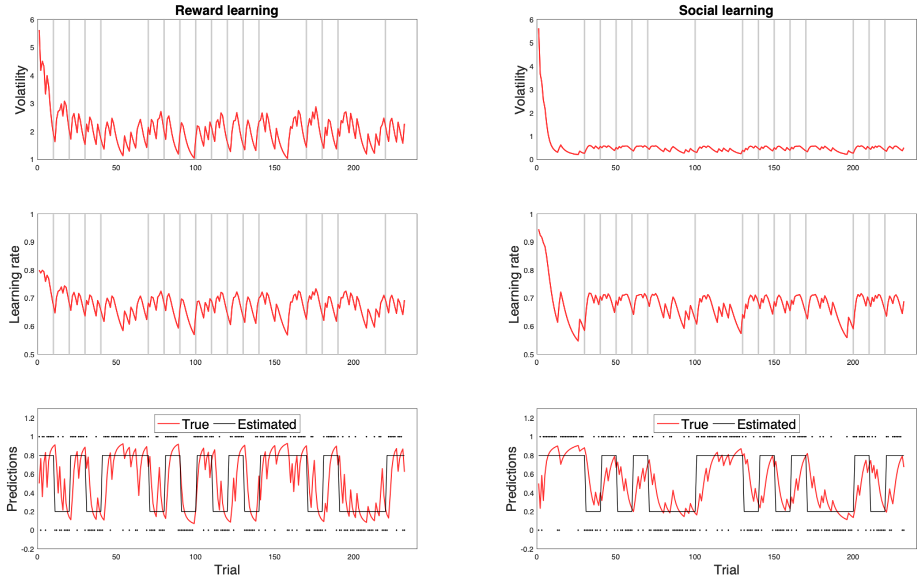


Supplementary Figure 3. An example of learning trajectory (subject S27) with parameter estimates $v_0 = 5.6$, $\lambda_r = 0.47$, $\omega_r = 1.4$, $\lambda_s = 0.50$, $\omega_s = 0.33$. Volatility, learning rate, and prediction signals for reward learning (left) and social learning (right).

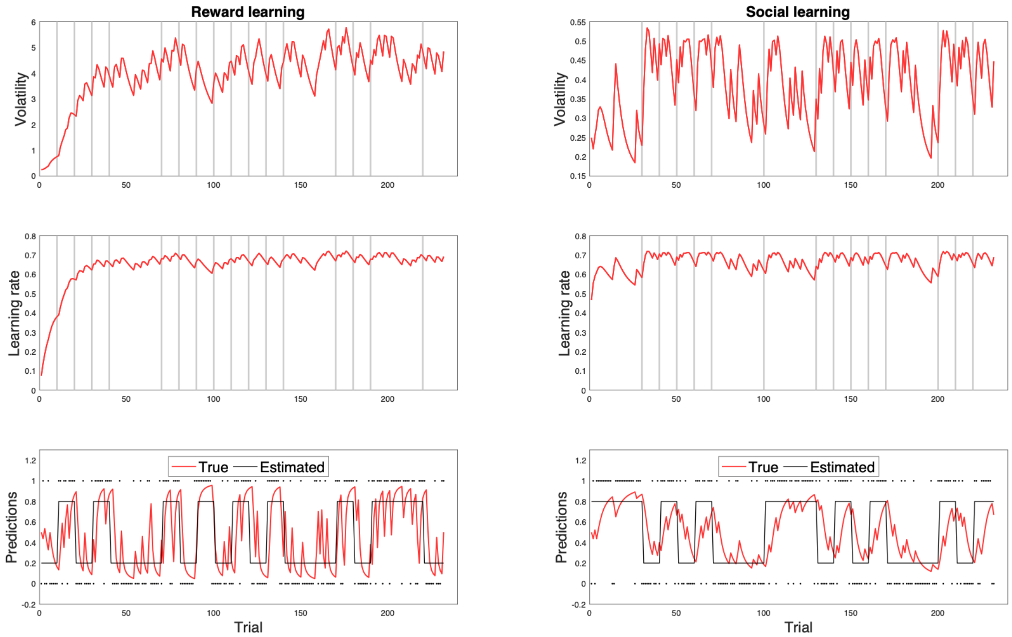


Supplementary Figure 4. Another example of learning trajectory (subject S30) with parameter estimates $v_0 = 0.25$, $\lambda_r = 0.21$, $\omega_r = 3.1$, $\lambda_s = 0.53$, $\omega_s = 0.29$. Volatility, learning rate, and prediction signals for reward learning (left) and social learning (right).

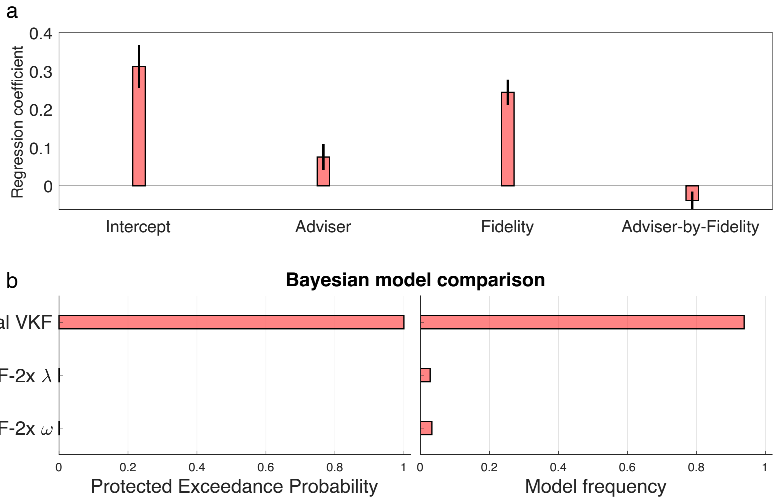


Supplementary Figure 5. Analysis of influences of advisers on learning fidelity. a) A logistic regression analysis with four regressors was conducted, which examines effects of advice-following (i.e. intercept), differential effects in following advisers (appetitive minus aversive), fidelity of the adviser on previous trial, and interaction of adviser and fidelity. Unlike the first three regressors that each showed a significant influence on choice, the interaction effect was not significant. b) Bayesian model comparison revealed that the original model with one set of noise parameters for learning was better than two alternative models that contained different noise parameters.

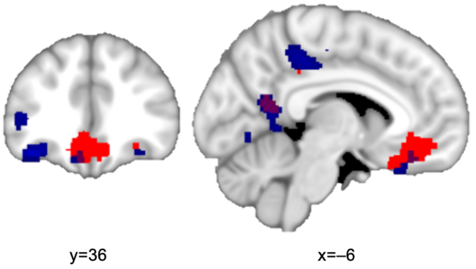


Supplementary Figure 6. Decision related activity: correlation with experience-based probability (red) and social-based probability (blue) of the chosen option. These activities overlap in the VMPFC (P<0.001 uncorrected).

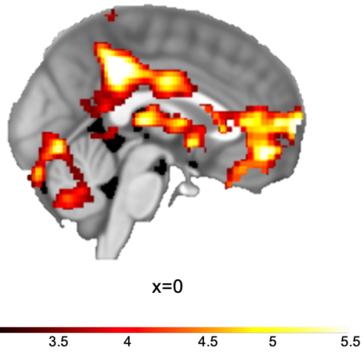


Supplementary Figure 7. Experience-based prediction error activity (P<0.001 uncorrected for illustration). Similar to previous studies with similar probabilistic learning tasks (Piray et al., 2019b), we found significant effects in the striatum extending into the amygdala, the ventromedial PFC, and the posterior cingulate.

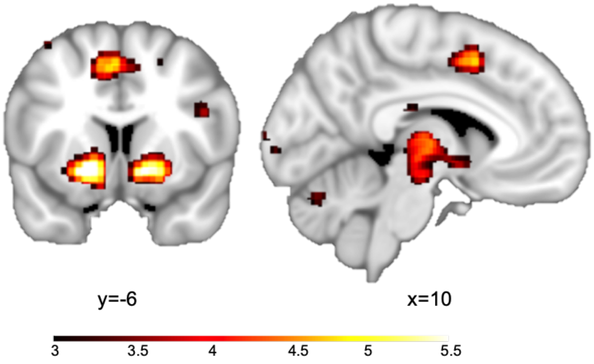


Supplementary Figure 8. Experience-based volatility activity (P<0.001 uncorrected for illustration). Similar to previous studies (Piray et al., 2019b), this effect is significant in the dorsal anterior cingulate cortex (peak at x=–10, y=6, z=48, voxel-level familywise small-volume corrected at p<0.05). There was also significant volatility related activity in the ventral striatum.

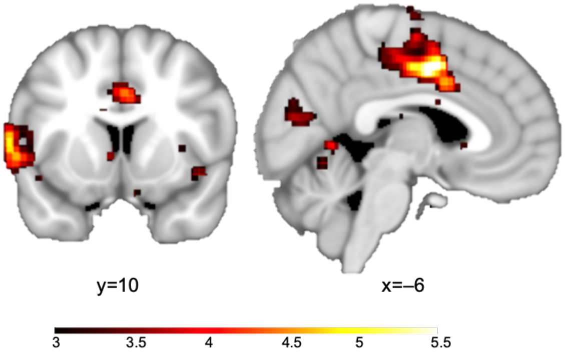


Supplementary Figure 9. Social volatility activity (P<0.001 uncorrected for illustration). We found significant deactivation in the midcingulate cortex and the temporal pole extending to TPJ.

| | $v_0$ | $\lambda_r$ | $\omega_r$ | $\lambda_s$ | $\omega_s$ | $\beta$ | $b$ | $w_{ap}$ | $w_{av}$ |
|---|---|---|---|---|---|---|---|---|---|
| Hierarchical fit | 4.13 | 0.05 | 5.11 | 0.69 | 0.51 | 1.29 | 1.19 | 0.60 | 0.69 |
| Individual fit | 11.59 | 0.27 | 5.52 | 0.50 | 1.86 | 1.34 | 1.16 | 0.60 | 0.67 |

Supplementary Table 1. Fitted parameters of the winning model for the hierarchical Bayesian inference used for model comparison and individual fit (in which the winning model was fitted to each participant's data separately) used for statistical analyses. Average parameters across all subjects.

| | $v_0$ | $\lambda_r$ | $\omega_r$ | $\lambda_s$ | $\omega_s$ | $\beta$ | $b$ | $w_{ap}$ | $w_{av}$ |
|---|---|---|---|---|---|---|---|---|---|
| Hierarchical fit | 1.82 | 0.05 | 5.28 | 0.62 | 0.3 | 1.39 | 0.95 | 0.64 | 0.64 |

Supplementary Table 2. Parameters used for generating regressors for fMRI analysis based on a variant of M1, which has equal weight parameters for both advisers (i.e. $w_{ap} = w_{av}$).
